# Nitrogen and phosphorus deficiencies alter primary and secondary metabolites of soybean roots

**DOI:** 10.1101/2022.03.14.484309

**Authors:** Mahnaz Nezamivand-Chegini, Sabine Metzger, Ali Moghadam, Ahmad Tahmasebi, Anna Koprivova, Saeid Eshghi, Manijeh Mohammadi-Dehchesmeh, Stanislav Kopriva, Ali Niazi, Esmaeil Ebrahimie

## Abstract

Nitrogen (N) and phosphorus (P) are two essential plant macronutrients that can limit plant growth by different mechanisms. We aimed to shed light on how soybean respond to low nitrogen (LN), low phosphorus (LP) and their combined deficiency (LNP). Generally, these conditions triggered changes in gene expression of the same processes, including cell wall organization, defense response, response to oxidative stress, and photosynthesis, however, response was different in each condition. A typical primary response to LN and LP was detected also in soybean, i.e., the enhanced uptake of N and P, respectively, by upregulation of genes for the corresponding transporters. The regulation of genes involved in cell wall organization showed that in LP roots tended to produce more casparian strip, in LN more secondary wall biosynthesis occurred, and in LNP reduction in expression of genes involved in secondary wall production accompanied by cell wall loosening was observed. Flavonoid biosynthesis also showed distinct pattern of regulation in different conditions: more anthocyanin production in LP, and more isoflavonoid production in LN and LNP, which we confirmed also on the metabolite level. Interestingly, in soybean the nutrient deficiencies reduced defense response by lowering expression of genes involved in defense response, suggesting a role of N and P nutrition in plant disease resistance. In conclusion, we provide detailed information on how LN, LP, and LNP affect different processes in soybean roots on the molecular and physiological levels.

## Introduction

Plant growth and development are largely determined by 17 macro/micro-nutrients (Pandey et al., 2021). Nitrogen (N) and phosphorus (P) are two macro-nutrients involved in many different biological processes and often limiting plant performance (Jiang et al., 2007; Nacry et al., 2013). Nitrogen limitation mostly affects enzyme activities that are required for energy metabolism such as photosynthesis and respiration (De Groot et al., 2003), while phosphorus limitation influences rather structural compounds (phospholipids and nucleic acids) (Marschner, 2011). Both deficiencies lead to numerous molecular and developmental adaptations (López-Bucio et al., 2003).

Besides structural roles of N and P, nitrate and phosphate function as signaling molecules to regulate gene expression and activate nutritional responses (Hu et al., 2019). It has been reported that their signaling pathways are not completely independent and they interact at several levels (Bloom et al., 1985; Chapin et al., 1987; Danger et al., 2008; Krouk & Kiba, 2020; Torres-Rodríguez et al., 2021). Phosphate limitation causes reduction of uptake, translocation, and assimilation of nitrogen (De Magalhães et al., 1998; Gniazdowska & Rychter, 2000; Rufty Jr et al., 1990), while changes in nitrate availability can trigger phosphate starvation response pathway (PSR) even in the phosphate sufficiency condition (Hu et al., 2019; Krouk & Kiba, 2020). One identified mechanism in N P crosstalk is through NIGT/HHO proteins that in high nitrate repress the expression of AtSPX1, AtSPX2, and AtSPX4, which in turn releases activity of PHR1 transcription factor and induce the phosphate starvation response (PSR) pathway (Kiba et al., 2018; Ueda et al., 2020).

To date, there have been a large number of studies on plant adaptation mechanisms to N and P deficiencies (Ishikawa et al., 2018; X. Liu et al., 2020; Ma & Chen, 2021; Zeng et al., 2018; H. Zhang et al., 2017). N deficiency triggers different responses in plants, such as reduction in photosynthesis and metabolism rate, or altering root architecture, which in turn increases the nitrogen absorption capacity (Han et al., 2014; M. Liu et al., 2020), improving N assimilation (Bi et al., 2005), and nodule formation in legumes for rhizobia symbiosis to utilize the atmospheric N_2_ (Harper, 1974). Phosphorus limitation is also associated with remodeling of root morphology (Zhao et al., 2004), increasing exudation of organic acids and acid phosphatase (Jiang et al., 2003), and formation of symbiotic interaction with mycorrhiza fungi (Liu et al., 2008) and rhizobia (Qin et al., 2011). Meanwhile, the combined deficiency of N and P have been studied in a few plant species. It was reported that low N can suppress P starvation response and in NP deficiency the low N response dominates (Hu et al., 2019; Krouk & Kiba, 2020; Torres-Rodríguez et al., 2021).

As soybeans contain a high concentration of protein, they require relatively larger amounts of N and P than cereal crops (Li et al., 2011; Ohyama et al., 2013). Soybean, as a legume, has gained specific characteristics during the plant evolution that help them to cope with N deficiency. One of them is nodule formation to utilize the atmospheric N_2_ through a molecular conversation with rhizobia (González & Marketon, 2003), the second one is isoflavonoid biosynthesis. Isoflavonoids are legume-specific group of phenylpropanoids that are involved in N_2_-fixing symbiosis and pathogen inhibition (Ferguson & Mathesius, 2003; Phillips & Kapulnik, 1995). The first enzyme in phenylpropanoid pathway is phenylalanine ammonia-lyase (PAL) that is the link between primary and secondary metabolism. The biosynthesis of flavonoids is committed by chalcone synthase (CHS) that catalyzes the iterative condensation of one p-coumaroyl CoA and three malonyl-CoAs to produce naringenin chalcone or isoliquiritigenin chalcone by cooperation with chalcone reductase (CHR) (Deng & Lu, 2017; Jia et al., 2015). The latter enzyme is mainly found in legumes (Bomati et al., 2005). In the next step, chalcone isomerase catalyzes the cyclization of these two compounds to naringenin flavanone or isoliquiritigenin flavanone (Ngaki et al., 2012). While isoliquiritigenin flavanone is a precursor for isoflavonoids (genistein, daidzein, and glycitein) in legumes or aurones in some specific species, naringenin can lead to different branches of flavonoids, including flavonols (kaempferol, quercetin, myricetin, isorhamnetin), flavandiols, anthocyanins (cyanidine, delphinidin, and petunidin), proanthocyanidins, and also isoflavonoids (Deng & Lu, 2017). Next steps in synthesis of flavonols, anthocyanins, and proanthocyanidins are catalyzed by flavanone 3-hydroxylase (F3H), flavonic 3’-hydroxylase (F3’H), or flavonoid 3’,5’-hydroxylase (F3’5’H). On the other hand, flavone synthase (FNS) convert flavanones to flavones and isoflavone synthase (IFS) is the key enzyme to redirect naringenin or liquiritigenin to isoflavonoid production (Deng & Lu, 2017). Several transcription factors (TFs) have been reported to regulate the production of different downstream branches from flavanons, including WD40, WRKY, BZIP, MADS-box, and R2R3-MYB (Ramsay & Glover, 2005; Stracke et al., 2007). Also, in soybean several TFs from MYB family have been identified as positive or negative regulators of some key genes in isoflavonoid biosynthesis (Bian et al., 2018; Chu et al., 2017; Han et al., 2017; Li et al., 2013; Yi et al., 2010). Interestingly, one specific TF can have dual roles for in the same gene family involved in flavonoid biosynthesis. For example, GmMYB176 is a positive regulator for some members of CHS gene family and a negative regulator for other members (Anguraj Vadivel et al., 2019).

In this study, we surveyed the effects of N, P, and combined NP deficiencies on the transcriptome profile of soybean roots. Additionally, as genes involved in flavonoid biosynthesis pathways were significantly enriched in these conditions, we measured flavonoid profiles in root and shoot. In addition, we explored gene expression pattern involving in cell wall reprogramming. Our results provided new hints as to which branches of flavonoid pathway may be activated and how root architecture might change during N and P deficiencies.

## Material and methods

### Plant growth condition

Soybean seeds (*Glycine max* cv. Williams 82.) were sterilized by sodium hypochlorite solution (1%, 5 min) and hydrogen peroxide (2%, 30 min). Sterilized seeds were planted in 40 pots containing river sand (4 seeds per 1-liter pot) in a completely randomized design with three replicates. Based on the experimental plan, pots were watered with 4 different solutions which were nitrogen normal-phosphorus normal (control), nitrogen normal-phosphorus deficiency (LP), nitrogen deficiency-phosphorus normal (LN), and nitrogen deficiency-phosphorus deficiency (LNP). The composition of nutrient solutions is listed in Supplemental Table S1 (Chiasson et al., 2014). NH_4_NO_3_ was used as N source in concentration 2.5 mM and 0.5 mM for control and deficiency, respectively, while for P, KH_2_PO_4_ and K_2_HPO_4_ were used at concentrations of 120 and 30 μM for normal condition and 20 and 5 μM for deficient condition. To inhibit nodulation, all solutions were prepared with sterile water, and also the sand was autoclaved at 121 °C for 15 minutes. The pots were watered with the nutrient solution once in 2-days for 28 days after planting. For sampling, three pots were randomly selected for each treatment, and roots of two seedlings in each pot were harvested, carefully cleaned and pooled and frozen in liquid N_2_ and stored at -80 °C for further molecular analysis. The rest of seedlings were harvested for physiological and morphological analyses.

### Assessments of plant growth parameters and nutrient content

To investigate effects of N and P deficiency on morphology and physiology of soybean, plant growth parameters (such as length, dry and fresh weight of root and shoot) were determined. The inorganic anions nitrate, phosphate, and sulfate were measured by ion chromatography (IC) as described by Dietzen et al. in 2020. The weighed and dried leaf and root tissues were extracted in 1.0 ml of pure Milli-Q water and incubated for 60 minutes at 4 °C, while shaking at 1500 rpm. Afterwards, the samples were transferred to 95 °C for 15 minutes. To remove the debris, they were centrifuged at 4 °C for 1 h at 12000 rpm. 100 μl of the supernatants were diluted with 900 μl pure Milli-Q water and measured with the Dionex ICS-1100 chromatography system and separated on a Dionex IonPac AS22 RFIC 4x 250 mm analytic column (Thermo Scientific, Darmstadt, Germany). 4.5 mM Na_2_CO_3_/1.4 mM NaHCO_3_ was used as running buffer. Standard curves were generated by using the external standards of 0.5 mM, 1 mM, and 2 mM KNO_3_, K_2_HPO_4_, and K_2_SO_4_. The mineral composition of leaves and roots was determined by inductively coupled plasma mass spectrometry (ICP-MS) by the CEPLAS Plant Metabolism and Metabolomics Laboratory, University of Cologne. 100–200 mg, fresh plant material was dried overnight in the oven at 60 C, digested in 0.5 ml of 67% nitric acid at 100°C for 30 min and diluted 1:10 in Milli-Q water. Samples were analyzed using an Agilent 7700 ICP-MS (Agilent Technologies, Waldbronn, Germany) following the manufacturer’s instructions. All reported IC and ICP data are means of 3-5 biologically independent samples.

### Flavonoid profiling

Flavonoids were extracted using 100 μl of extraction buffer (5% hydrochloric acid in methanol, plus 50 nM of D3-Sakuranetin as internal standard) per 10 mg of fresh plant material. Samples were sonicated for 15 min following extraction and then hydrolyzed at 100 °C for 10 min to obtain the aglycones. Samples were centrifuged at 15,000g. The LC-MS analysis was performed using an Agilent 1260 HPLC System (Agilent, Waldbronn, Germany) connected to a QTRAP 5500 mass spectrometer equipped with a Turbo V Source (Sciex, Darmstadt, Germany). The chromatographic separation was performed on a Kinetex C_18_ column (2.6 μm, 100 Å, 4.6 mm x 50 mm, Aschaffenburg, Germany). A linear gradient elution of eluents A (0.01% formic acid) and B (methanol, 0.01% formic acid) starting with the following elution program 0 – 0.5 min 5% B, 0.5-3 100% B, 3-6 min 100% B, 6-6.1 min 5 % B, 6.1 - 10 min 5 % B, was used for the separation. The flow rate was 0.5 ml/min, the injection volume was 10 μl. The detection and quantification was accomplished in the positive mode using multiple-reaction monitoring (MRM). The raw data were analyzed using Analyst 1.6 software (Sciex, Darmstadt, Germany). Integrated peak areas from the inbuilt Analyst Quantitation Wizard were manually corrected. Peak areas of each compound for each sample were normalized to the internal standard, and quantification was performed by external calibration to standards (naringenin, kaempferol, quercetin, isorhamnetin, cyaniding, delphinidin, genistein, apigenin, daidzein and glycitein from Sigma-Aldrich; Darmstadt, Germany).

### RNA isolation, RNA sequencing, and qRT-PCR

Total RNA was isolated from the frozen roots of soybean seedlings with Trizol (Invitrogen, Carlshad, CA) following the manufacturer’s protocol. RNA quantity and quality were determined using a Nanodrop2000 Spectrophotometer (Thermo Scientific), and by agarose gel electrophoresis. Further three step, including assessment of RNA integrity (using an Agilent Technologies 2100 Bioanalyzer), cDNA library construction (TrueSeq Standard mRNA LT Sample Prep Kit) and sequencing (100 bp paired reads using Illumina.NovaSeq6000) were done by Macrogen company (Korea).

For RT-qPCR analysis, DNase treatment to remove contamination of genomic DNA and first-strand cDNA synthesis were performed using QuantiTect Reverse Transcription Kit (Qiagen). Quantitative real-time RT-PCR (qPCR) was performed for some genes involved in phenylpropanoid biosynthesis and phosphate-deficiency marker genes, using gene-specific primers (Supplemental Table S2) as in (Koprivova et al., 2019). The Actin-6 (*GLYMA_18G290800*) gene was used as the internal reference gene for qRT-PCR because its transcript levels were not affected by the treatments in the RNAseq data.

### Read preprocessing and mapping

From the 12 samples which were sent to Macrogen Inc, only 8 passed quality control based on their RINs, including all 3 replicates of control, 2 replicates of LP, 1 replicate of LN, and 2 replicates of LNP. From these 8 samples 100 bp paired-end reads were generated. An initial quality control analysis was performed on FastQ files using FastQC software (Andrews, 2010), then preprocessing of raw sequence data was conducted and low quality reads, adapter sequences and duplicate mapping reads were filtered using Trimmomatic on linux. The preprocessed FastQ files were aligned to soybean reference genome using STAR. The output of this alignment was stored in a file format called SAM. In order to quantify gene expression level of this combined N and P experiment, the count values obtained from STAR outputs were used to form count matrix as input file for DESeq2 R package (Love et al., 2014). The count matrix of 8 samples was used for detection differential expressed genes in three conditions in comparison with normal condition using Dexus package (Klambauer et al., 2013). Dexus is a statistical model using a Bayesian framework by an expectation-maximum (EM) algorithm that does not need replicates to detect differentially expressed genes (DEGs). We used its supervised method with default parameters. The method provides an threshold for informative/non-informative (I/NI) values to extract DEGs (Klambauer et al., 2013).

### GO, KEGG enrichment, and data analysis

The GO functional classification and KEGG pathway analyses were conducted by the DAVID (Database for Annotation, Visualization and Integrated Discovery) software (Huang et al., 2009) for three DEG lists obtained from Dexus. The scatter plots of top terms in GO and KEGG results were made by using ggplot2 R package (Wickham, 2011). Visualization of flavonoid and isoflavonoid KEGG pathways were constructed using fold changes of genes annotated in these pathway by Pathview online tool (Luo et al., 2017).

### Statistical analysis

All statistical analyses were performed using R software packages. The experimental data were subjected to statistical Tukey test and analysis of variance (ANOVA). Correlation coefficients for ionome data and correlation network for metabolome and transcriptome data were calculated by Hmisc package and visualized by ggcorrplot and Cytoscape, respectively.

## Results

### The effect of N, P, and the combined NP deficiency on root and shoot biomass

Nitrogen and phosphorus deficiency caused changes in soybean root and shoot biomass. Generally, LN and LNP showed the same trends; in the deficient conditions shoot length, shoot fresh weight, root dry weight, shoot dry weight, and root to shoot dry weight ratio showed significant changes in comparison with soybean plants grown in the normal condition. However, only differences in root dry weight and root to shoot dry weight ratios were significant in LP (Table 1). Shoot length, shoot fresh weight, and shoot dry weight were significantly decreased in LN and LNP, while shoot traits did not change in LP in comparison with the normal condition. On the other hand, root dry weight and root to shoot dry weight have significantly increased in LN, LP, and LNP compared to the normal conditions. Considering no significant changes in root length in all three conditions but increase in root dry weight, it might be concluded that soybean root developed more lateral roots to increase the root surface for N and P uptake. In our results, shoot was less sensitive to P deficiency but more sensitive to low N.

**Table 1.** Mineral nutrient concentrations in mg/kg DW of soybean root and leaf under nitrogen (LN), phosphorus (LP), and the combined deficiency (LNP). Mean values of 3 biological replicates are shown. The significant differences (Tukey test <0.05) are indicated with asterisks with different superscript letters.

| elements | Root |  |  |  | leaf |  |  |  |
| --- | --- | --- | --- | --- | --- | --- | --- | --- |
|  | ctrl | LN | LP | LNP | ctrl | LN | LP | LNP |
| Mg | 2168.5<sup>b</sup> | 2613<sup>a</sup> | 1729.1<sup>c</sup> | 2431.3<sup>a</sup> | 2232<sup>a</sup> | 2260.3<sup>a</sup> | 2220.2<sup>a</sup> | 2327.7<sup>a</sup> |
| P | 1866.2<sup>a</sup> | 1670.83<sup>b</sup> | 1213.8<sup>c</sup> | 1517<sup>b</sup> | 1923.8<sup>a</sup> | 2006<sup>a</sup> | 1685.8<sup>a</sup> | 1697.1<sup>a</sup> |
| S | 4678.5<sup>a</sup> | 4836.5<sup>a</sup> | 3652.5<sup>b</sup> | 4676.1<sup>a</sup> | 3681.5<sup>ab</sup> | 4192.5<sup>b</sup> | 2648<sup>c</sup> | 3467.1<sup>b</sup> |
| K | 6885.6<sup>b</sup> | 8051<sup>a</sup> | 4741.3<sup>c</sup> | 6417.3<sup>b</sup> | 12716.3<sup>b</sup> | 14977.7<sup>a</sup> | 10189.4<sup>c</sup> | 12021<sup>b</sup> |
| Ca | 13673.7<sup>b</sup> | 13620.9<sup>b</sup> | 11149.3<sup>c</sup> | 14337.2<sup>a</sup> | 20463.6<sup>b</sup> | 18030.3<sup>c</sup> | 18481.6<sup>bc</sup> | 22627.9<sup>a</sup> |
| Cr | 3.2<sup>a</sup> | 3.8<sup>a</sup> | 3.6<sup>a</sup> | 3.8<sup>a</sup> | 2.1<sup>c</sup> | 2.3<sup>b</sup> | 2.4<sup>a</sup> | 2.4<sup>a</sup> |
| Mn | 72<sup>b</sup> | 81.3<sup>b</sup> | 69.2<sup>b</sup> | 156<sup>a</sup> | 56<sup>b</sup> | 51.5<sup>b</sup> | 53.2<sup>b</sup> | 75.3<sup>ab</sup> |
| Fe | 862.2<sup>b</sup> | 733.5<sup>b</sup> | 593.2<sup>b</sup> | 2080.9<sup>a</sup> | 147.2<sup>a</sup> | 124.4<sup>b</sup> | 126.2<sup>b</sup> | 114.2<sup>b</sup> |
| Cu | 25.3<sup>a</sup> | 12<sup>c</sup> | 18.2<sup>b</sup> | 13<sup>c</sup> | 10.6<sup>a</sup> | 9<sup>b</sup> | 9.2<sup>ab</sup> | 9<sup>b</sup> |
| Zn | 119.8<sup>a</sup> | 103.5<sup>b</sup> | 88<sup>c</sup> | 100.9<sup>b</sup> | 83.3<sup>a</sup> | 77.6<sup>a</sup> | 91.7<sup>a</sup> | 76<sup>a</sup> |
| As | 1<sup>ab</sup> | 0.7<sup>bc</sup> | 0.5<sup>c</sup> | 1.2<sup>a</sup> | 0.2<sup>a</sup> | 0.1<sup>b</sup> | 0.1<sup>c</sup> | 0.1<sup>c</sup> |
| Sr | 42.4<sup>ab</sup> | 44.6<sup>ab</sup> | 38.9<sup>b</sup> | 48.7<sup>a</sup> | 37.5<sup>a</sup> | 32.8<sup>a</sup> | 34.6<sup>a</sup> | 37<sup>a</sup> |
| V | 1.01<sup>b</sup> | 1.2<sup>b</sup> | 1.1<sup>b</sup> | 2.2<sup>a</sup> | 0.04<sup>a</sup> | 0.02<sup>c</sup> | 0.03<sup>b</sup> | 0.02<sup>c</sup> |
| Cd | 0.9<sup>a</sup> | 0.8<sup>a</sup> | 0.8<sup>a</sup> | 0.8<sup>a</sup> | 0.2<sup>a</sup> | 0.1<sup>c</sup> | 0.1<sup>b</sup> | 0.1<sup>b</sup> |
| Ni | 4.7<sup>c</sup> | 6<sup>b</sup> | 4.4<sup>c</sup> | 7.3<sup>a</sup> | 3<sup>a</sup> | 3.5<sup>a</sup> | 3.2<sup>a</sup> | 3.2<sup>a</sup> |
| Co | 2.1<sup>b</sup> | 1.7<sup>c</sup> | 2.5<sup>a</sup> | 2.4<sup>ab</sup> | 0.2<sup>a</sup> | 0.1<sup>b</sup> | 0.2<sup>a</sup> | 0.2<sup>a</sup> |

### Anion and elemental content of soybean under N, P, and the combined NP deficiency

Next, we conducted IC and ICP-MS analyses to determine how the deficiencies affected the elemental contents and anion concentrations in the root and leaves of soybean (Table 2 and Figure 1, respectively). Nitrate was under detection limit in leaves, however, it showed a remarkable decrease in root under all three deficiencies compared with normal condition (Figure 1). Sulfate and phosphate concentrations did not change in both leaf and root under LP condition. Nonetheless, sulfate showed the same pattern in LN and LNP conditions; increasing in root and decreasing in leaf. Phosphate increased in LN in root and leaf, but decreased in LNP root.

**Table 2.** Mineral nutrient concentrations in mg/kg DW of soybean root and leaf under nitrogen (LN), phosphorus (LP), and the combined deficiency (LNP). Mean values of 3 biological replicates are shown. The significant differences (Tukey test <0.05) are indicated with asterisks with different superscript letters.

| Root |  |  |  |  | leaf |  |  |  |
| --- | --- | --- | --- | --- | --- | --- | --- | --- |
| elements | ctrl | LN | LP | LNP | ctrl | LN | LP | LNP |
| Mg | 2168.5<sup>b</sup> | 2613<sup>a</sup> | 1729.1<sup>c</sup> | 2431.3<sup>a</sup> | 2232<sup>a</sup> | 2260.3<sup>a</sup> | 2220.2<sup>a</sup> | 2327.7<sup>a</sup> |
| P | 1866.2<sup>a</sup> | 1670.83<sup>b</sup> | 1213.8<sup>c</sup> | 1517<sup>b</sup> | 1923.8<sup>a</sup> | 2006<sup>a</sup> | 1685.8<sup>a</sup> | 1697.1<sup>a</sup> |
| S | 4678.5<sup>a</sup> | 4836.5<sup>a</sup> | 3652.5<sup>b</sup> | 4676.1<sup>a</sup> | 3681.5<sup>ab</sup> | 4192.5<sup>b</sup> | 2648<sup>c</sup> | 3467.1<sup>b</sup> |
| K | 6885.6<sup>b</sup> | 8051<sup>a</sup> | 4741.3<sup>c</sup> | 6417.3<sup>b</sup> | 12716.3<sup>b</sup> | 14977.7<sup>a</sup> | 10189.4<sup>c</sup> | 12021<sup>b</sup> |
| Ca | 13673.7<sup>b</sup> | 13620.9<sup>b</sup> | 11149.3<sup>c</sup> | 14337.2<sup>a</sup> | 20463.6<sup>b</sup> | 18030.3<sup>c</sup> | 18481.6<sup>bc</sup> | 22627.9<sup>a</sup> |
| Cr | 3.2<sup>a</sup> | 3.8<sup>a</sup> | 3.6<sup>a</sup> | 3.8<sup>a</sup> | 2.1<sup>c</sup> | 2.3<sup>b</sup> | 2.4<sup>a</sup> | 2.4<sup>a</sup> |
| Mn | 72<sup>b</sup> | 81.3<sup>b</sup> | 69.2<sup>b</sup> | 156<sup>a</sup> | 56<sup>b</sup> | 51.5<sup>b</sup> | 53.2<sup>b</sup> | 75.3<sup>ab</sup> |
| Fe | 862.2<sup>b</sup> | 733.5<sup>b</sup> | 593.2<sup>b</sup> | 2080.9<sup>a</sup> | 147.2<sup>a</sup> | 124.4<sup>b</sup> | 126.2<sup>b</sup> | 114.2<sup>b</sup> |
| Cu | 25.3<sup>a</sup> | 12<sup>c</sup> | 18.2<sup>b</sup> | 13<sup>c</sup> | 10.6<sup>a</sup> | 9<sup>b</sup> | 9.2<sup>ab</sup> | 9<sup>b</sup> |
| Zn | 119.8<sup>a</sup> | 103.5<sup>b</sup> | 88<sup>c</sup> | 100.9<sup>b</sup> | 83.3<sup>a</sup> | 77.6<sup>a</sup> | 91.7<sup>a</sup> | 76<sup>a</sup> |
| As | 1<sup>ab</sup> | 0.7<sup>bc</sup> | 0.5<sup>c</sup> | 1.2<sup>a</sup> | 0.2<sup>a</sup> | 0.1<sup>b</sup> | 0.1<sup>c</sup> | 0.1<sup>c</sup> |
| Sr | 42.4<sup>ab</sup> | 44.6<sup>ab</sup> | 38.9<sup>b</sup> | 48.7<sup>a</sup> | 37.5<sup>a</sup> | 32.8<sup>a</sup> | 34.6<sup>a</sup> | 37<sup>a</sup> |
| V | 1.01<sup>b</sup> | 1.2<sup>b</sup> | 1.1<sup>b</sup> | 2.2<sup>a</sup> | 0.04<sup>a</sup> | 0.02<sup>c</sup> | 0.03<sup>b</sup> | 0.02<sup>c</sup> |
| Cd | 0.9<sup>a</sup> | 0.8<sup>a</sup> | 0.8<sup>a</sup> | 0.8<sup>a</sup> | 0.2<sup>a</sup> | 0.1<sup>c</sup> | 0.1<sup>b</sup> | 0.1<sup>b</sup> |
| Ni | 4.7<sup>c</sup> | 6<sup>b</sup> | 4.4<sup>c</sup> | 7.3<sup>a</sup> | 3<sup>a</sup> | 3.5<sup>a</sup> | 3.2<sup>a</sup> | 3.2<sup>a</sup> |
| Co | 2.1<sup>b</sup> | 1.7<sup>c</sup> | 2.5<sup>a</sup> | 2.4<sup>ab</sup> | 0.2<sup>a</sup> | 0.1<sup>b</sup> | 0.2<sup>a</sup> | 0.2<sup>a</sup> |

**Figure 1.**
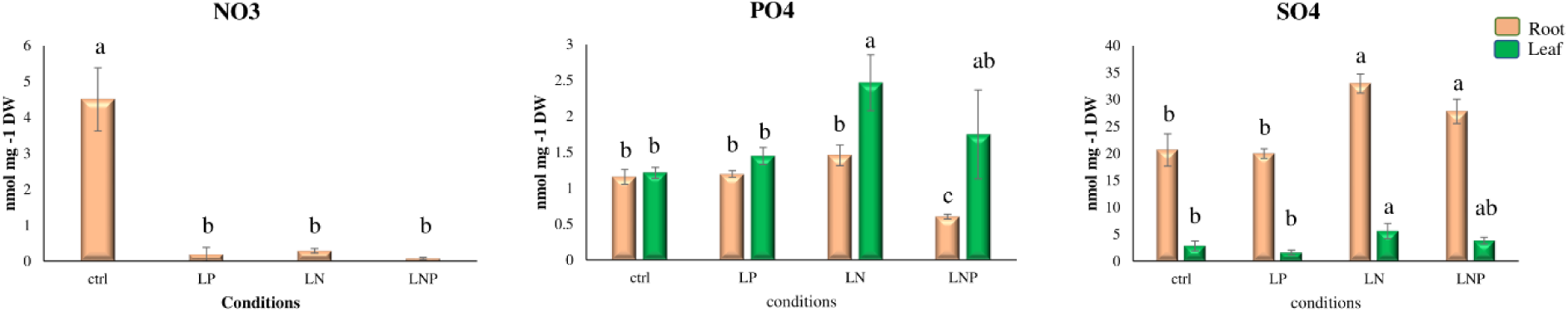
Anion accumulations in soybean root and leaf under LN, LP, and LNP conditions. Data are given as the mean ± standard deviation for root and leaf in orange and green bars, respectively. The significant differences (Tukey test < 0.05) are indicated with asterisks with different superscript letters.

Since phosphate levels were not affected in LP, we measured total P and other elemental contents to test whether the LP condition in our experiment was sufficient. The ICP-MS data showed that both leaf and root contained less P in LP in comparison with control, but the decrease in leaf was not statistically significant due to high biological variation. Also total S content was lower in LP compared to control, although we did not observe significant changes of sulfate in LP. It seems, in our P deficiency, total P and S were more significantly affected than their anion forms, pointing unexpectedly to more significant effects on the organic P and S pools. Among other nutrients, potassium (K) showed an opposite trend in LN and LP; soybean under LP had less K in both root and leaf compared with normal condition, but both tissues under LN condition stored more K than in control. Interestingly, the LNP plants showed very significant alterations in the ionome, in the roots they possessed ca. two-fold higher concentration of iron, manganese, nickel, and vanadium than the controls, while copper and zinc contents were lower. In the leaves, Fe was significantly lower abundant, as well as Cu, while Zn content was increased (Table 2).

### RNA sequencing data analysis and identification of Differentially Expressed Genes (DEG)

To obtain insights into the molecular mechanisms of the response of soybean to N and P deficiencies, RNA was isolated from roots and subjected to RNA sequencing. Eight sample libraries, from RNAs with a sufficient RIN, were sequenced and the summary of their sequencing data is shown in Supplemental Table S3. Over 550 million reads were obtained across these eight libraries, ranging from 63 to 79 million reads from each (Supplemental Table S3). After quality control of reads and removal of adaptor contamination, reads were mapped to the soybean reference genome.

Since we did not have three biological replications of the RNAseq data for all treatments, we used a Dexus package, which allows to detect differentially expressed genes (DEGs) even without biological replicates (Klambauer et al., 2013). The differentially expressed genes (DEGs) with a cut off I/NI value more than 0.1, which represents a high differential expression threshold corresponding to specificity > 99 % according the Dexus calculation, and P-value of 0.05 in response to N, P, and the combined deficiency were detected. In each condition, different numbers of genes were found to be differentially regulated in root in comparison with normal conditions (Figure 2A). In response to P starvation, 2482 DEGs were obtained, 950 genes were upregulated and 1532 genes downregulated (Figure 2A, Supplementary file). The corresponding numbers for LN was 2429 genes, including 809 upregulated genes and 1620 downregulated ones. The number of DEGs in response to combination of both factors was 1955 with 343 genes upregulated and 872 genes downregulated (Figure 2A).

**Figure 2.**
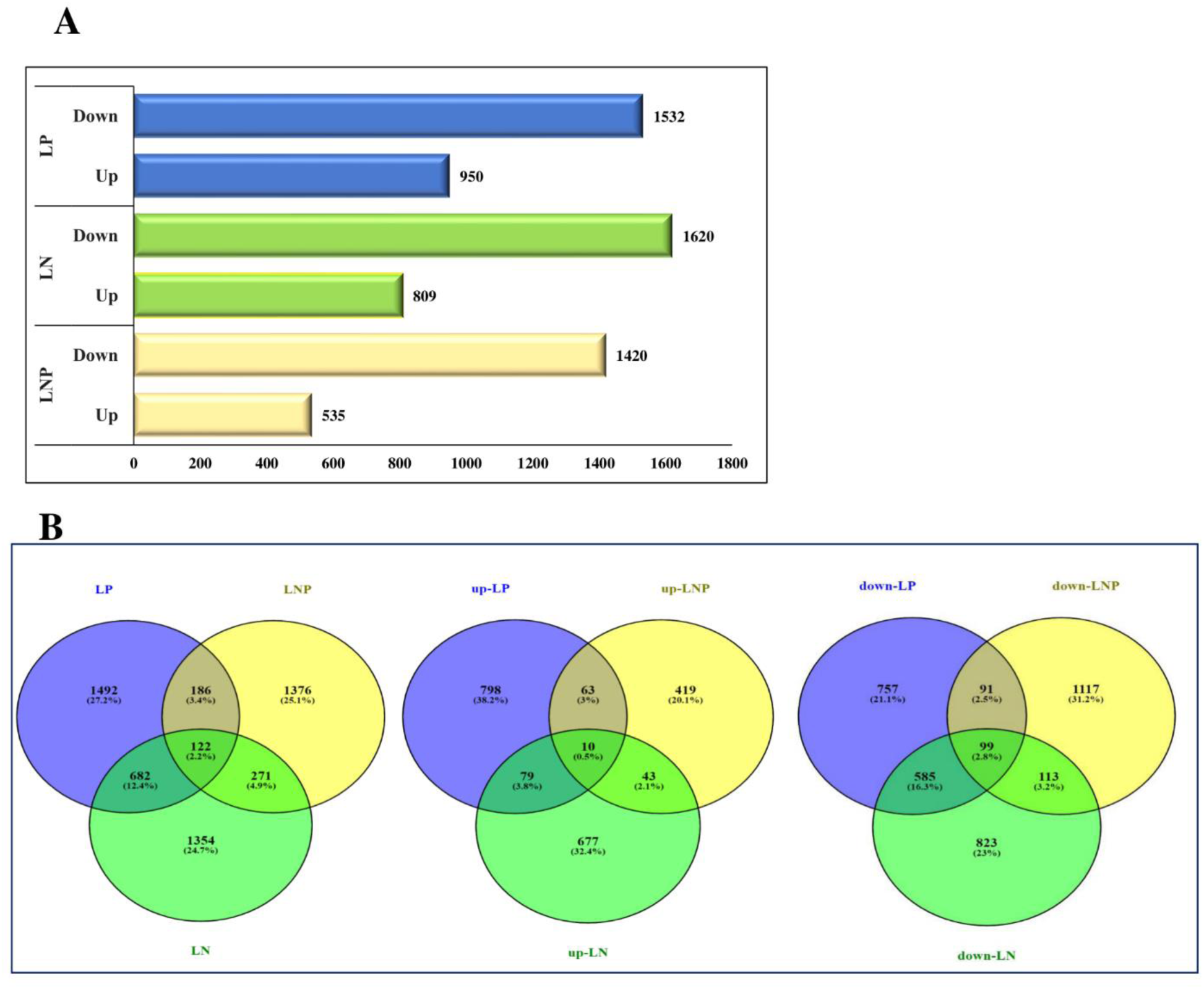
A. The number of up and down-regulated DEGs in nitrogen, phosphorus, and the combined deficiency conditions. B. Venn diagrams showing overlaps for total DEGs, upregulated and downregulated DEGs in the three conditions of LN, LP, and LNP.

The overlap of DEGs in the three pair-wise comparisons with normal condition are illustrated in Venn diagram in Figure 2B. In total, 122 genes were found to be differentially regulated by all three conditions (Supplementary file). Interestingly, 99 were down-regulated in all three conditions, including 16 NB-ARC domain containing proteins and 11 TIR domain containing proteins, which are involved in defense response signaling and cell death programming with increasing of ROS production (Song et al., 2021). In addition, 10 genes were upregulated at all three conditions and 13 genes were regulated in a different way in the individual deficiencies. Four genes involved in C metabolism, fructose-1,6 bisphosphatase, ribulose bisphosphate carboxylase/oxygenase activase, Ribulose bisphosphate carboxylase small chain 1, as well as one chlorophyll a-b binding protein were upregulated only in LN but downregulated in the two other conditions. On the other hand, one sugar transporter, sugar efflux transporter *SWEET15* was upregulated in LP, in line with previously reported link between P deficiency and upregulation of sugar transporters (Xue et al., 2018).

To confirm the gene expression patterns detected in the transcriptome dataset and also to check the marker genes for phosphate and nitrate deficiency, we selected three genes to analyze by qRT-PCR: *PHT1*.*1, PHT1*.*13*, and NRT1/PTR family *6*.*1*. PHT1.1 and PHT1.13 as members of PHT1 transporter family are responsible for phosphate uptake and translocation (Nussaume et al., 2011). The selected member of NRT/PTR family was commonly downregulated in all conditions. Transporters of NRT1/PTR family function as low-affinity transporters (Sakuraba et al., 2021) and show down-regulation in response to N deficiency in plants (Islam, 2019; Lejay et al., 1999) and P starvation in barley (Long et al., 2019). The transcript levels of these three genes obtained by qRT-PCR were consistent with the patterns apparent in the RNAseq analysis; *PHT1*.*1* and *PHT1*.*13* were upregulated in LP, while NRT/PTR family 6.1 showed significant reduction in transcript levels in LN and LNP (Supplemental Figure S1A). We also used expression data of SPX domain-containing proteins in these conditions, described in (Nezamivand-Chegini et al., 2021). Also these expression data showed a high correlation (r^2^= 0.93) between qRT-PCR and RNAseq with 6 genes upregulated by P deficiency (Supplemental Figure S1B).

### GO and KEGG enrichment analysis of DEG

To gain a better understanding of the DEGs function, GO functional enrichment of DEGs affected in LN, LP, and LNP was conducted. For this analysis, the DEGs in each three sets were used and GO terms with adjusted *P*-values < 0.05 were considered significantly enriched (Supplementary file). The top GO terms in each category; biological processes (BP), molecular functions (MF), and cellular components (CC) for each conditions are shown in Figure 3. Among BP, “protein-chromophore linkage”, “photosynthesis”, “response to oxidative stress”, and “plant-type secondary cell wall biogenesis” were the most enriched terms in all three conditions. However, genes involved in photosynthesis and plant-type cell wall organization were differentially regulated in different conditions. Another GO term enriched in all conditions was “plant-type secondary cell wall biogenesis. LP conditions induced expansins, pectinesterases, and casparian strip membrane proteins. Interestingly, genes for casparian strip synthesis were not induced in LN and LNP (Supplemental Table S4). Instead, in LN genes for cellulose synthase A (CESA) catalytic subunits, many expansins, and xyloglucan endotransglucosylase/hydolases (XTH) were upregulated, and putative pectinesterase inhibitors, most of xyloglucan endotransglucosylase/hydrolases, and callose synthases were downregulated. In LNP, on the other hand, all (CESA) catalytic subunits, COBRA proteins, pectin acetylestrases, putative beta-1,4-xylosyltransferases, putative galacturonosytransferases, putative pectinesterase inhibitors, and putative xyloglucan endotransglucosylase/hydrolases were downregulated and only one expansin and pectinestrase showed a slight induction.

**Figure 3.**
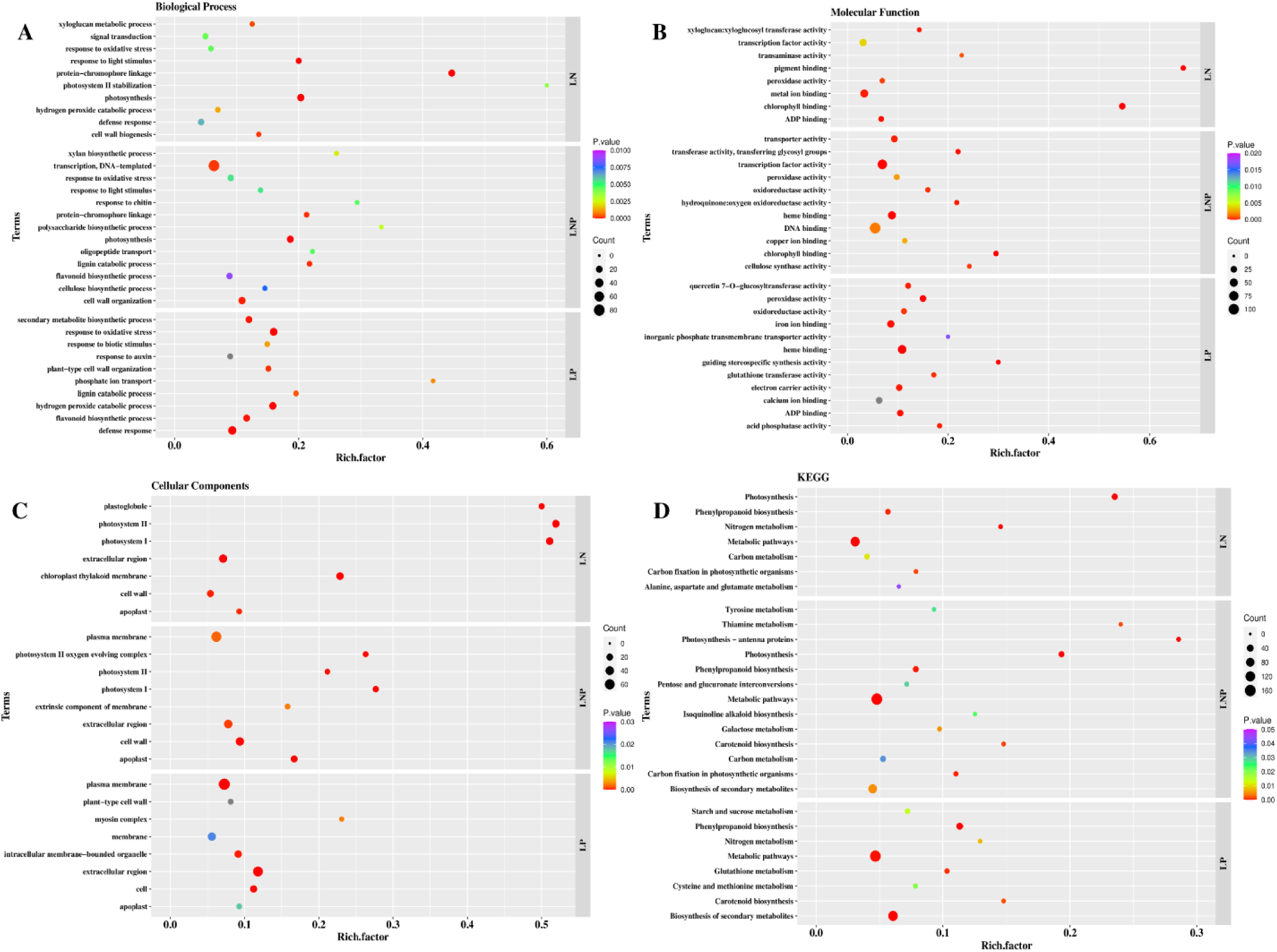
GO and KEGG functional enrichment of DEGs corresponding to LN, LP, and LNP conditions. The Y-axis represent terms and X-axis indicates the ratio of the differentially expressed gene number to the total gene number for the term (Rich factor). Significance of the GO term in the indicated gene list is represented by the point fill color. The number of DEGs in the gene list that are annotated to each term is represented by point size. A to C represent three branches of GO; biological processes (BP), Molecular functions (MF), and Cellular Components (CC), respectively and D represents KEGG pathways.

To further understand the physiological processes involved in responses to N and P deficiency, enrichment of KEGG pathways was analyzed (Figure 3 D). The most enriched pathways in LN and LP was “biosynthesis of secondary metabolites”, whereas in LN condition “Alanine, asparate and glutamate metabolism” (Supplementary file). In the LN, additionally, DEGs in the processes related to nitrogen metabolism and carbon metabolism were enriched, confirming the link between nitrogen acquisition and photosynthetic activity as reported before (Coruzzi & Zhou, 2001; Curci et al., 2017; Ranathunge et al., 2014; S.-y. Yang et al., 2015). Also under LN six high affinity nitrate transporters were upregulated, but downregulation of NRT1, glutamate synthase (GOGAT), and glutamate dehydrogenase (GDH), suggested increase in uptake capacity for nitrate but reduction in utilization of N for protein synthesis.

### Transcriptome and metabolome profiles of flavonoids under LN, LP, and LNP conditions

Since legumes are characteristic by the synthesis of isoflavonoids, we asked whether these metabolites might play a role in the response to N and P deficiency. To this end, we used pathview to visualize the fold-changes of soybean genes annotated in flavonoid biosynthesis pathway (00941) and isoflavonoid biosynthesis pathway (00943) in LN, LP, and LNP conditions (Figure 4). The expression pattern of genes from flavonoid pathway of soybean roots was different in LP in comparison with the two other conditions, especially in the regulation of *F3’H*, anthocyanidin synthase (*ANS)*, and anthocyanidin reductase 1 (*ANR1*). F3’H converts naringenin or dihydrokaempferol to dihydroquercetin, then dihydroquercetin could be converted to anthocyanidins by two reactions catalyzed by dihydroflavonol reductase (DFR) and ANS (Petrussa et al., 2013), but the synthesis of anthocyanidins could branch off the anthocyanin pathway to catechin production by enzymatic activity of ANR (Bogs et al., 2005). Furthermore, it seems that in LP, soybean root transcriptome is modulated to produce anthocyanins, flavonols, and catechin, while in LN and LNP these pathways were down-regulated and probably other flavonoid branches might be up-regulated. Indeed, genes involved in isoflavonoid biosynthesis were up-regulated in LN and LNP, but down-regulated in LP (Figure 4).

**Figure 4.**
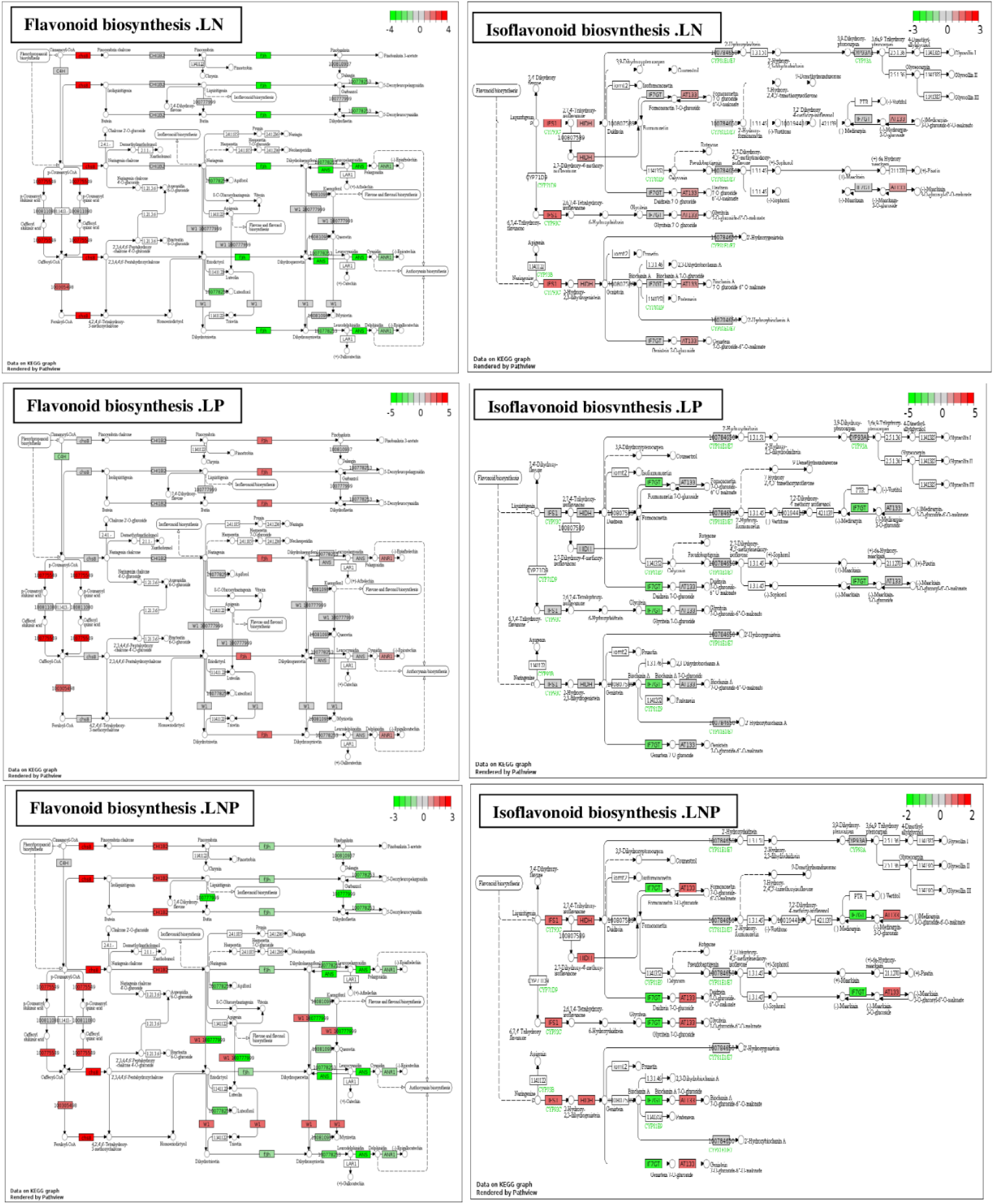
Soybean gene expression pathway analysis of Flavonoid biosynthesis (left) and isoflavonoid (right) under LN, LP, LNP conditions top-down, respectively.

To test whether the modulation of transcripts of genes from the flavonoid pathways results in alteration of the metabolite levels, the flavonoid profiles of root and shoot extracts were obtained by LC-MS analysis (Figure 5). The flavonoid profiles of shoot and root were different from those reported previously (Deng et al., 2019; Ho et al., 2002) since we could not detect any anthocyanins in roots, whereas in the shoots, cyanidin and delphinidin were present in all 4 conditions. Furthermore, we detected flavanone (naringenin), flavone (apigenin), flavonols (kaempferol, quercetin, and isorhamnetin), and isoflavonoids (genistein, daidzein, and glycitein) in both tissues. Except daidzein and glycitein, all other compounds accumulated in shoots to higher levels than in roots (Figure 5). Among flavonols, kaempferol levels decreased in all deficiencies in both roots and shoot, whereas quercetin content was reduced in NLP. In shoots quercetin was reduced also in LP, whereas in roots it was lower in LN that in control (Figure 5). Isorhamnetin accumulated to higher levels in root in both LN and LP, but only in LN in shoot, where it was slightly reduced in LP. Naringenin showed reduced accumulation in root in LN and LNP, and in shoot in LP. Consistently with enrichment of the isoflavonoid biosynthesis pathway in the RNAseq data, we observed a significant increase of glycitein in both root and shoot in LN and LNP, and significant increase of genistein and daidzein in LNP condition in root and shoot, respectively. Despite the expression of anthocyanin synthesis genes in the roots, we could not detect any anthocyanin in this tissue, however, we observed increased accumulation of two anthocyanins, cyanidin and delphinidin, in shoots in LP condition. Possibly, anthocyanin produced in root might be translocated to shoot.

**Figure 5.**
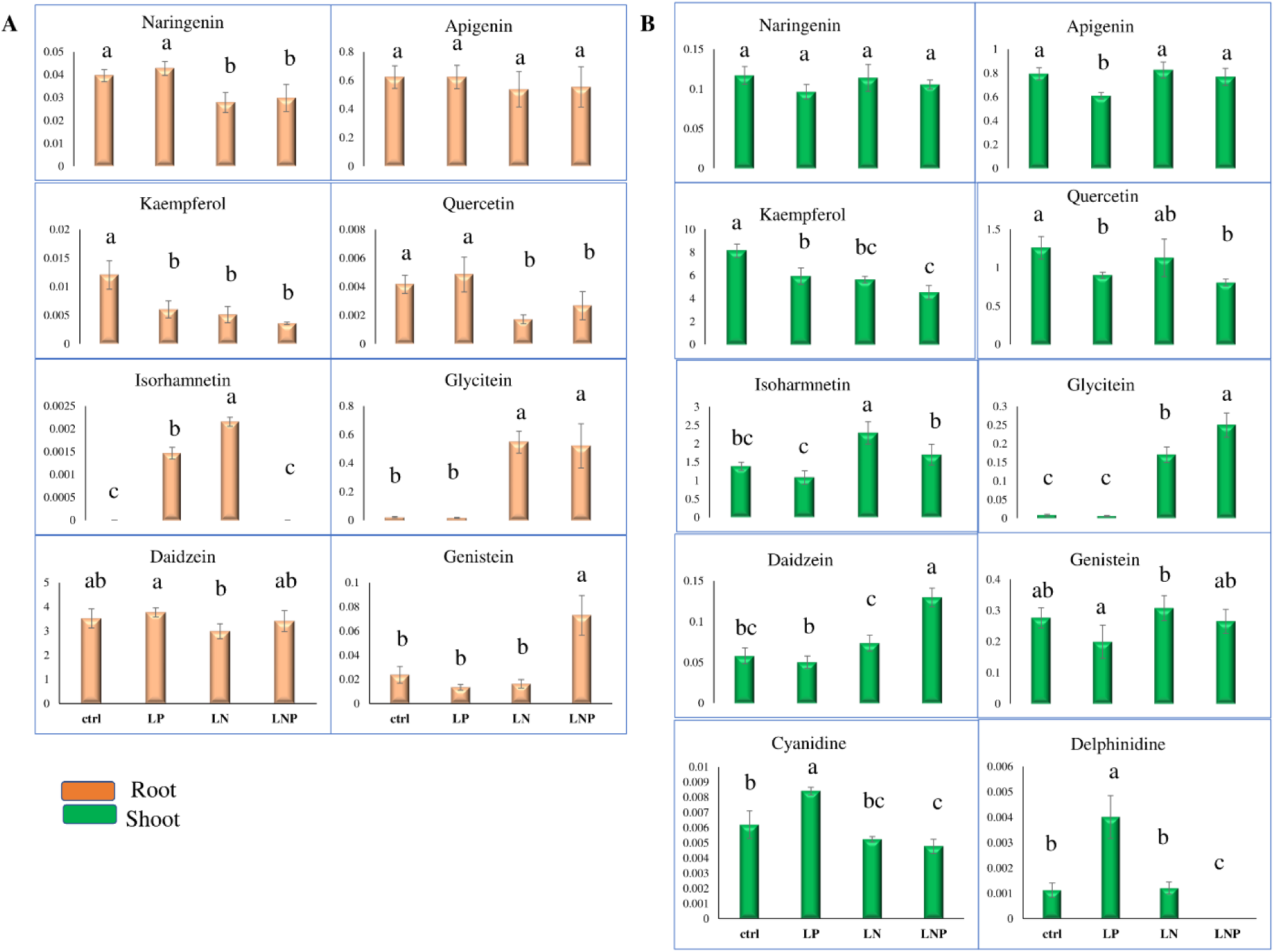
Flavonoid accumulation in soybean root (A) and shoot (B) in control and low nitrogen (LN), low phosphorus (LP), and the combined deficiency (LNP). Data are shown as peak area and as means ± SD. The significant differences (Tukey test <0.05) are indicated with asterisks with different superscript letters.

To investigate the correlation of metabolites and gene expression, we calculated correlations using RPKMs from the RNAseq analysis and the obtained concentrations of flavonoids and isoflavonoids from LC-MS analysis. The constructed network with 73 nodes and 283 edges was visualized using Cytoscape (Figure 6). At metabolite level, quercetin (0.95), genistein (−0.74), and daidzein (0.83) showed correlation to naringenin, the first compound starting flavonoid and isoflavonoid biosynthesis. Daidzein also showed positive correlation to apigenin (0.83) and quercetin (0.78). As for correlation between genes and metabolites, 22 genes correlated with the flavonoid compounds, mostly genes involved in the first steps of phenylpropanoid pathway, such as shikimate O-hydroxycinnamoyltransferase, spermidine hydroxycinnamoyl transferase, trans-cinnamate 4-monooxygenase, and stemmadenin O-acetyltransferase, for the first compounds of both flavonoid and isoflavonoid biosynthesis. Genistein and kaempferol negatively correlated with genes in anthocyanin production such as *ANR1* and *ANS*. Some members of chalcone synthase gene family also showed correlation with metabolites; *CHS13* was correlated with apigenin (0.9) and daidzein (0.71), but *CHS10* was negatively correlated to kaempferol (−0.71). *F3H*, the gene involved in both anthocyanin and flavonol production, showed negative correlation with genistein (−0.8), but positive correlations with daidzein (0.76), quercetin (0.83), and naringenin (0.83).

**Figure 6.**
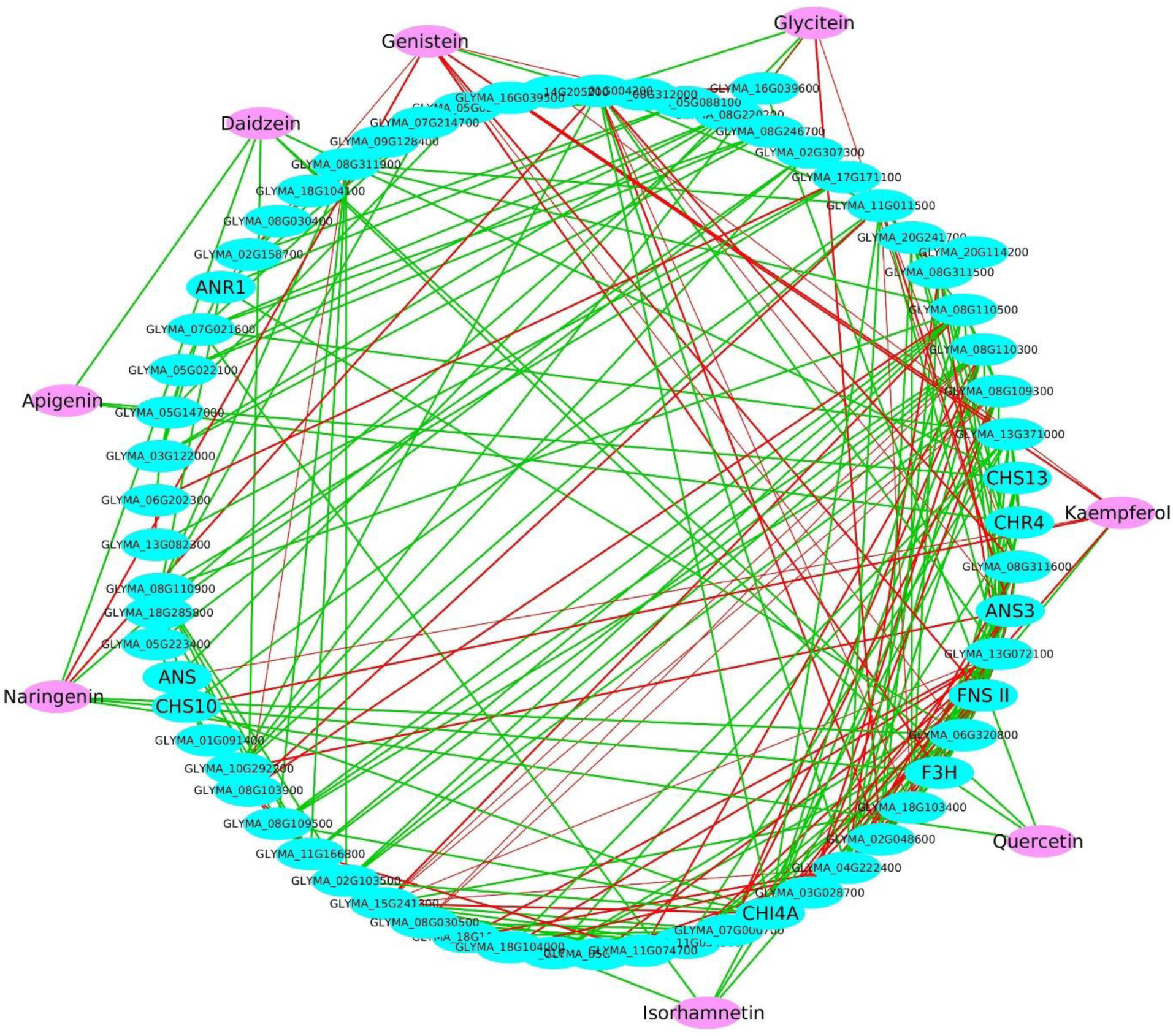
The constructed correlation network using metabolite and RNA-seq data from soybean root. The network was constructed by isoflavonoid and flavonoid compound (pink) and genes annotated in these two pathway (blue color).

**Figure 8.**
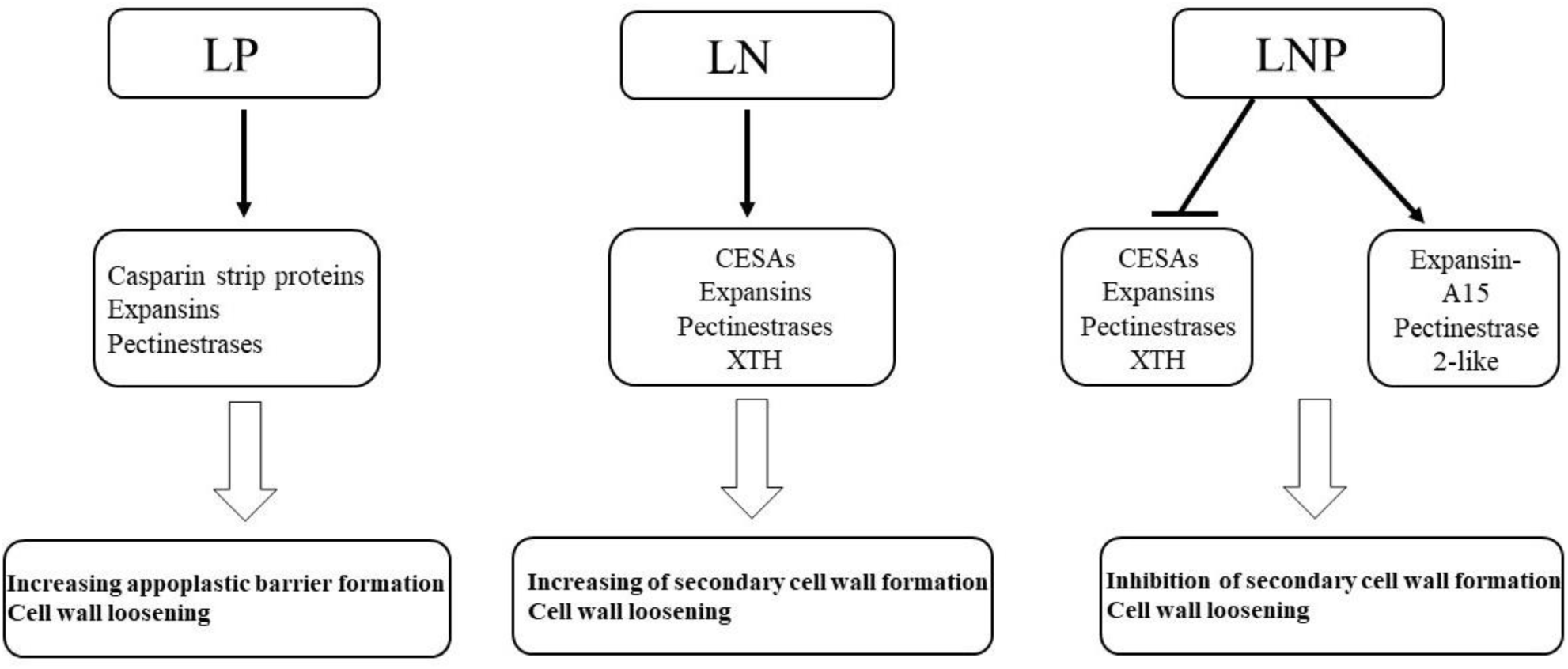
Processes underlying cell wall reprogramming in soybean roots in response to LN, LP, and LNP conditions.

In legumes, especially in soybean, the regulation pattern of flavonoid genes might be more complex than in other species, as the key genes are mostly encoded by gene families not a single genes and contain an extra branch of flavonoid metabolism (García-Calderón et al., 2020). To verify and extend the RNAseq data, we determined transcript levels of four key genes involved in phenylpropanoid pathway by RT-qPCR (Supplemental Figure S2). The gene for PAL1, the first enzyme starting phenylpropanoid biosynthesis, showed a significant increase only in response to LN in root. Transcript levels of *CHS8*, a member of CHS gene family involved in diverting naringenin chalcone to flavonoid and isoflavonoid branches showed more than two-fold increase in all three deficiencies compared to the normal condition. This might suggest that these two pathway branches are induced in the deficiency conditions, consistently with the results from RNAseq and flavonoid metabolite measurement.

## Discussion

N and P are key macronutrients limiting plant growth and development since plants require greater amounts of N than of any other element and due to the low mobility of P in soil (Yoneyama et al., 2012), (Heuer et al., 2017). Nitrate and phosphate are two main available forms of N and P that also function as signal molecules (Hu et al., 2019), having large impacts on transcriptome, proteome, metabolome, and ionome composition. So far, substantial progress in our understanding of the individual pathways of nitrate and phosphate signaling have been achieved, but their cross-talk is poorly understood. An antagonistic interaction between Pi and nitrate has been discussed for legumes (Ma & Chen, 2021); Pi limitation causes increase in Pi uptake, but decrease in nitrate uptake and inhibition of nodule formation (Sulieman & Tran, 2015). On the other hand, sufficient N induces PSR, even during high availability of phosphate. NIGT1 is one important key player in N P crosstalk; in LP condition and N-sufficiency, NIGT1 is induced and promotes Pi uptake, but reduces nitrate influx and accumulation (Wang et al., 2020). Here, we descibe how non-symbiotic soybean reacts in LN, LP, nad LNP at transcriptome, metabolome, and ionome levels.

Transcriptome analysis resulted in almost the same number of DEGs in response to LN and LP and slightly fewer DEGs in LNP condition. In each condition genes known to be involved in mechanisms of response to N and P deficiencies were induced. For example, the increase in organic acids and purple acid phosphatase and upregulation of phosphate transporters are two well-documented strategies in LP condition (Delhaize et al., 2009; Liang et al., 2012). Xue et al (2018) reported upregulation of several PHT1 transporters and *purple acid phosphatase* (*PAPs*) in soybean nodules (Xue et al., 2018). Consistently, in our study several PHT1 and PAPs were upregulated to enhance Pi mobilization and acquisition in LP. Also, SPX genes involved in PSR were differentially expressed in LP condition. *GmSPX1/10* are orthologous to *AtSPX3* and *GmPHO1;1* is orthologus of *AtPHO1;H1* (Nezamivand-Chegini et al., 2021), which are involved in regulation of phosphate homeostasis (Duan et al., 2008). Here, we obtained the same results and proved the conclusion that some SPX genes are regulated in the opposite pattern in N and P deficiency, in fact, LN could suppress PSR in the LNP conditions. To exemplify, GmSPX10 was upregulated under LP condition in both RNAseq and RT-qPCR, but downregulated in LNP. Interestingly, several NB-ARC domain containing proteins and 11 TIR domain containing proteins were significantly down regulated in LN and LP, which points to a link between N and P deficiency and reduction in plant defense activity. As for low N, soybean roots depicted a distinct trend to improve nitrogen use efficiency (NUE) under LN. NUE, could be divided into N uptake efficiency (NUpE) and N utilization efficiency (NUtE) (Moll et al., 1982). Our results showed that at LN some high affinity nitrate transporters were upregulated, while glutamate synthase (GOGAT) and glutamate dehydrogenase (GDH) were downregulated in root. It shows that N management was mainly affected by N absorption by upregulation of high affinity nitrate transporters and thus the gene regulation in LN had more impact on improvement of NUpE.

There is link between NUtE and the inhibition of photosythetic capacity in N deficienct conditions. Contribution of NUpE and NUtE in NUE varies among different species and varieties (Barraclough et al., 2010; Gaju et al., 2011). For example, comparing two different cultivars of wheat, synthetic hexaploid wheat-derived (SDCs) and nonsynthetic-derived cultivars (NSCs), SDCs showed higher NUtE and NUE than NSCs and lower inhibition of photosynthetic capacity under N deficiency in leaf (M. Liu et al., 2020). Generally, chlorophyll content in LN and LP exhibits an opposite pattern; decrease in LN but increase in LP (Bassi et al., 2018; Bechtaoui et al., 2021). The increase in chlorophyll content in LP is due to the lack of Pi molecules to sustain the photosynthesis, particularly, RUBISCO and fructose-1,6-bisphosphatase function (Bechtaoui et al., 2021). Consistently, these two genes were downregulated in both LP and LNP, but upregulated in LN in our data. Altogether, we could conclude that in leaf, N deficiency decreased the photosynthetic capacity. With the concominant upregulation in NUpE genes and downregulation in NUtE genes, soybean leaves might show reduction in chlorophyll content but excess in P content, as photosynthesis and N assimilation, two energy-dependent processes are affected. However, soybean might cope with the N deficiency pthrough the help of roots. Although roots are heterotrophic organs (Aschan & Pfanz, 2003), chloroplast accumulation can be observed in upper parts of primary root near hypocotyl junction (Kobayashi et al., 2012). Therefore, expression of genes involved in photosynthesis in the root can give insight into regulation of predominantly leaf processes, as in rice photosynthesis related genes were regulated by N starvation to the same extent in roots and leaves (W. Yang et al., 2015). Indeed, photosynthesis was an enriched term among DEGs common in LN and LNP conditions (Figure 3). The DEGs for photosynthesis showed opposite expression pattern in responses to LN and LNP, 23 DEGs were down-regulated in LNP and 39 DEGs were up-regulated in LN condition (Supplemental Table S4). Upregulation of these genes, encoding components of photosynthetic electron transport chain, during N starvation was reported in previous study in rice (W. Yang et al., 2015), as well as their suppression in Pi-deficient root (Kang et al., 2014; Li et al., 2010; Wu et al., 2003). The opposite pattern in LN and LNP thus might reflect compensation of low photosynthesis in leaf and overcoming the Pi excess in LN, with the limitation of both N and P, leading to downregulation of photosynthetic genes in LNP.

Another GO term enriched in all conditions was “plant-type secondary cell wall biogenesis”. Cell wall reprogramming is an important strategy for plant adaptations to biotic and abiotic stress, root development, and embryogenesis (Landi & Esposito, 2017; Ogden et al., 2018). Cell walls are highly dynamic and mainly composed by polysaccharides, such as cellulose, pectins, and callose, as well as lignin and structural proteins (Fukuda, 2014; Ogden et al., 2018). Different enzymes are involved in cell wall reprogramming, such as expansins, xyloglucan endotransglucosylase/hydrolase, peroxidase, and enzymes involved in pectin modification (Franciosini et al., 2017; Tenhaken, 2015). Expansins have important roles in cell wall loosening and are often regulated by nutrient stress to subsequently affect plant growth and nutrient uptake (Muller et al., 2007). Pectins are linked with cellulose and they have important roles in cell wall remodeling. Pectin de-esterification causes stifenning of cell wall and hindrance of cell wall loosening by expansins (Hocq et al., 2017). In our RNAseq results, different genes, encoding cell wall biogenesis enzymes were regulated by the nutrient deficiencies. Intriguingly, we observed different enzymes being affected in LN, LP, and LNP conditions (Supplemental Table S4). LP conditions induced casparian strip membrane proteins, while in LN and LNP, CESA genes, expansins, and XTH genes were differentially regulated in comparison with normal condition. Casparian strip, made of suberin, functions as an apoplastic barrier to the diffusion of water and is often affected by nutrient availability (Schreiber et al., 1999). Indeed, K and S deficiency result in inreased suberization, while Mn, Fe, and Zn limitation leads to delay in suberization (Barberon et al., 2016; Ogden et al., 2018). Phosphorus deficiency also resulted in an increase in apoplastic barrier formation (Li et al., 2020), agreeing with our results. Additionally, we observed upregulation of some expansin genes in LP that normally cause cell wall loosening. Interestingly, GmEXLB1 was shown to have an important role in alteration of soybean root architecture (Kong et al., 2019). As cell wall organization can be different in each root zone, it is possible that in response to phosphorus deficiency soybean root directs cell wall remodeling to appoplastic barrier formation and cell wall loosening.

Cellulose is synthesized by a large multi-meric protein complexes, which contain CESA proteins as the catalytic units (Endler & Persson, 2011). Cellulose synthase complexes (CSC) are involved in the formation of both primary and secondary cell walls; among different subunits of CESA, CESA1, CESA3, CESA6, CESA2, CESA5, and CESA9 are active in the primary cell wall formation, but CESA4, CESA7, and CESA8 are associated with the secondary cell wall formation (Endler & Persson, 2011). XTHs are involved in the rearrangement of xyloglucans by cutting and rejoining of them, causing reversible or irreversible loosening cell walls to permit cell expansion (Fry et al., 1992). XTH is mostly present in tissues in which expansion has ceased (Catalá et al., 2001; O’Donoghue et al., 2001), indicating that it has a role in the formation of secondary walls of xylem and/or phloem cells (Bourquin et al., 2002). Our transcriptome data shows that in LN condition, soybean roots increase expression of genes for secondary wall formation and cell wall loosening, but in LNP, no distinct behaviour can be discerned except cell wall loosening and inhibition of secondary cell wall formation. The effects of N availability on secondary cell wall formation and its components have been reported for different plants, including rice (W. Zhang et al., 2017), maize (Sun et al., 2018), poplar (Pitre et al., 2007), sorghum (Rivai et al., 2021), and Eucalyptus (Camargo et al., 2014), but the effects are different in different plant species. In Eucalyptus, N deprivation led to increase in lignin in secondary cell wall, but in sorghum, it caused accumulation of several hemicelluloses. One reason for the alteration of cell wall in N deficiency could be an adaptation strategy to limitation of N-containing molecules, such as proteins and amono acids and reduction in demand for carbon skeletons for N assimilation, so, plants may accumulate surplus carbon as starch or utilize it for cell wall materials (Rivai et al., 2021). This is consistent with the reduction in expressions of genes for nitrogen metabolism, such as GDH and GOGAT in our results. Our transcriptomics data thus showed a different cell wall reprogramming in different conditions (Figure 6). It seems that by appoplastic barrier formation or secondary wall formation, soybean roots try to make a balance in absorbtion of P or N in LN and LP, respectively. However, in LNP, because of low photosynthesis and low energy, it reduces expression of almost all genes for cell wall synthesis to save photosynthesis products and energy. While these conclusion need further experiments to verify, the morphology of root and shoot supports this idea. We observed increase in root/shoot ratio but not in root length, meaning that root growth was prevented, but casparian strip and secondary cell wall formation were induced. As a result, root lenghts did not show any changes, but root weight was increased. Detailed analyses of root cell walls would be necessary to prove this conclusion.

The plant ionome is tightly regulated by uptake, translocation, storage, and remobilization as plants aim to maintain their nutrient homeostasis (Courbet et al., 2019). Multiple cross-talks between macro-and micronutrients have been described previously (Fan et al., 2021). The phenomenon of interactions between N and P and also with other nutrients has been recognized, but the underlying mechanisms have not been completely understood. We compared the correlation of total P contents with other nutrients. Given the alterations in expression of genes for secondary wall synthesis it is possible, that some of the changes in ionome are caused by changes in casparian strip and endodermal barrier (Barberon et al., 2016; Huang & Salt, 2016). Several mutants lacking the ability of casparian strip formation have been reporter to display multielement ionomic phenotypes (Hosmani et al., 2013; Kamiya et al., 2015). The way that casparian strip can affect ionomic composition is that it prevents mineral nutrients entering the stele from cortex and stops the leak of minerals from stele back to cortex (Geldner, 2013). Based on our results, Pi showed positive correlation with K, S, Ca, and Zn. In LP conditions, accumulation of these minerals were decreased in both root and leaf. As for micronutrients, almost all micronutrients accumulated in lower amount in LP compared to normal condition.

Given the unique feature of legumes, the synthesis of isoflavonoids, and the regulation of genes for flavonoids and isoflavonoids in our RNAseq dataset, we assayed flavonoid and isoflavonoid accumulation in response to different conditions on the metabolome level. All in all, obtained metabolome profile showed consistent results with the pattern that we inferred from gene expression data suggesting that soybean regulates different branches of phenylpropanoid pathway in a different way depending on the specific deficiency condition. Phenolic compounds influence different physiological processes in plants, such as seed germination, cell division, and synthesis of photosynthetic pigments (Tanase et al., 2019). Also, their synthesis is increased in response to environmental stresses helping to plant adaptations, through their antioxidant properties, defensive mechanisms, and signal transduction as well as in nutrient uptake and mobilization (Sharma et al., 2019). So far, the effects of various abiotic stresses on polyphenol biosynthesis, such as drought, salinity, heavy metals, temprature, UV radiation, and heavy metals, have been investigated (Ancillotti et al., 2015; Handa et al., 2019; Naikoo et al., 2019; Smirnov et al., 2015), in which generally antioxidanive properties of polyphenols helped plants to be able to scavenge free radicals (Schroeter et al., 2002). Phenylpropanoid biosynthesis is regulated at activities of some key-points enzymes in this pathway. PAL is the first enzyme onseting phenypropanoid biosynthesis in response to stresses, especially responding to nitrogen deficiency. In fact, in N deficiency, due to accumulation of photosynthetic assimilated and low N, the biosynthesis of carbon-based secondary metabolites such as phenylpropanoid are increased. (Deng et al., 2019). Especially in legumes, nitrogen deficiency induced flavonoids and isoflavonoids as signals to indece transcription of the genes involved in the biosynthesis of Nod factors (García-Calderón et al., 2020). Flavonol and anthocyanin accumulation in N and P deficiency, also have been reported. In P deficiency, anthocyanin and flavonol accumulation are induced in arabidopsis and tobacco, respectively (Jia et al., 2015; Lei et al., 2011; Li et al., 2014). Dihydroflavonols are common substrate for DFR and FLS enzymes that because FLS transcripts were increased in tobacco under P deficiency, caused increase in flavonol instead of anthocyanin (Jia et al., 2015). Flavonoid biosynthesis in legume crops, is more interesting due to their ability to isoflavonoid biosynthesis playing important roles as signal for nodulation (Tohge et al., 2017).

Sugiyama et al (2016) assayed tissue isoflavone contents and the secretion of isoflavones from soybean at various growth stages and nutrient conditions. they reported in LN condition, isoflavone content, while in LP isoflavone secretion were reduced and isoflavone content were not significantly changed (Sugiyama et al., 2016). Also, they claimed that in N deficiency. Genistein was highly secreted. The secretion of isoflavones are occurred by two dustinct pathways; ATP-dependent transport, especially genistein and secretion into apoplast followed by hydrolysis with ICHG (isoflavone conjugates hydrolyzing beta-glucosidase) (Sugiyama et al., 2007). Our results, consistantly, showed increasing in isoflavone contents in LN and LNP conditions, but not significant alteration in LP. Comparing root and shoot, root had higher rate of daidzein/genistein in comparisom with shoot. This was also observed by Sugiyama et al (2016). However, we observed significant increasing of glycetin in both root and shoot in LN rather than daidzein and genistein, could be due to genetic variant between cultivars. LN and LNP, indeed, showed accumulation of isoflavones, but in the opposite pattern for genistein content; increasing in LNP and decreasing in LN. the reason could be due to the discrepency in the secretion of genistein in two conditions, as the secretion for genistein is ATP-dependent, it be said that in LNP, genistein more accumulated rather than secreted.

## Conclusion

Our results showed soybean changed the primary and secondary metabolites production to adapt to the deficiency conditions. In low N, the high-affinity nitrate transporters were upregulated but genes involved in nitrogen metabolism, such as glutamate synthase (GOGAT) and glutamate dehydrogenase (GDH) were downregulated in root. It suggests an increase in uptake of N but reduction in its utilization for protein synthesis. As a consequence, carbon surplus that is not used for N metabolism, needs to be used or stored. The alteration in cell wall biosynthesis directing it to produce secondary cell wall could be a strategy to use such surplus carbon. However, soybean, still is enduring N limitation and the other strategy to provide N was icreasing of PAL expression to release N from amino acids. In LP, soybean upregulated high-affinity phosphate transporters to uptake more P from the soil. The other strategy in LP, was enhancement of appoplastic barrier production to prohibit leaking P back to soil and selectively uptake nutrients. In LNP, however, soybean showed neither secondary wall nor appoplastic barrier formation, as it was struggling with both low N and P, simultaneously. In contrast with LN, it showed downregulation in photosynthesis gene expression and cellulose synthase genes. However, LNP and LN showed almost same patterns in flavonoid biosynthesis pathway; both of them showed decreasing in anthocyanin production but increase isoflavonoid production that this result were consistent with metabolome measurement. Understanding the strategies of soybean to cope with nutrient deficiencies will be the key to breed new cultivars adapted to low fertiliser input.

## Acknowledgements

We acknowledge the Ministry of Science, Research and Technology of Iran and the University of Shiraz for the exchange scholarship to University of Cologne. We thank to Sabine Ambrosius (University of Cologne) for technical assistance with the ICP-MS measurements. The MS Platform and SM is supported by the Cluster of Excellence on Plant Sciences (CEPLAS). Research in SK’s lab is funded by the Deutsche Forschungsgemeinschaft (DFG) under Germany’s Excellence Strategy – EXC 2048/1 – project 390686111.

## Supplementary material

Supplementary file.xlsx contains all DEGs, overlapped DEGs, as well as GO and KEGG outputs Additional file.docx contains supplemental tables and figures

---

**Supplemental Table S1.** Nutrient solution for soybean growth in N and P deficiency

| Chemicals | Final Concentration | Concentration (deficiency) |
| --- | --- | --- |
| Macro elements |  |  |
| NH<sub>4</sub>NO<sub>3</sub> | 2.5 mM | 0.5 mM |
| MgSO<sub>4</sub>.7H<sub>2</sub>O | 500 µM |  |
| KH<sub>2</sub>PO<sub>4</sub> | 120 µM | 20 µM |
| K<sub>2</sub>HPO<sub>4</sub> | 30 µM | 5 µM |
| KCl | 250 µM |  |
| Fe-Na-EDTA | 100 µM |  |
| CaCl<sub>2</sub> | 250 µM |  |
| Trace elements |  |  |
| H<sub>3</sub>BO<sub>3</sub> | 46 µM |  |
| MnSO<sub>4</sub>.H<sub>2</sub>O | 8 µM |  |
| ZnSO<sub>4</sub>.7H<sub>2</sub>O | 8 µM |  |
| CuSO<sub>4</sub>.5H<sub>2</sub>O | 2 µM |  |
| Na<sub>2</sub>MoO<sub>4</sub>.2H<sub>2</sub>O |  |  |

**Supplemental Table S2.**
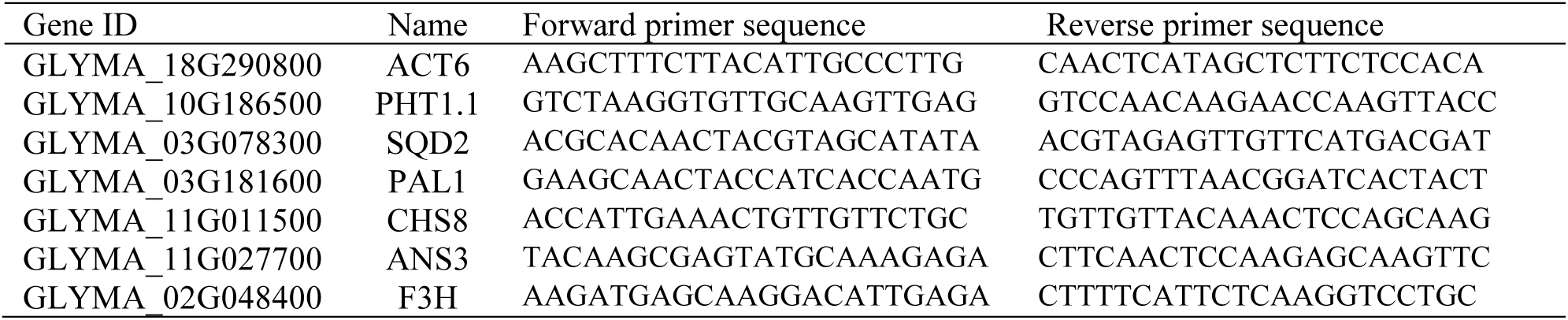
Primer sequences used for qRT-PCR

**Supplemental Table S3.** The summary of sequencing data.

| Sample ID | Total read (Mb) | GC content (%) | Q30 (%) | Uniquely mapped read (%) |
| --- | --- | --- | --- | --- |
| N1P1-1 | 65.2 | 45.5 | 94.3 | 94.5 |
| N1P1-2 | 70.6 | 45 | 94.3 | 94.01 |
| N1P1-3 | 71.9 | 45.2 | 93.4 | 92.71 |
| N1P0-1 | 78.4 | 45.5 | 93.7 | 93.71 |
| N1P0-2 | 63 | 45.7 | 94.2 | 93.3 |
| N0P1 | 79 | 45.6 | 93.8 | 94.56 |
| N0P0-1 | 65.4 | 44.7 | 94.1 | 94.56 |
| N0P0-2 | 64 | 45.7 | 93.5 | 93.22 |

**Supplemental Table S4.**
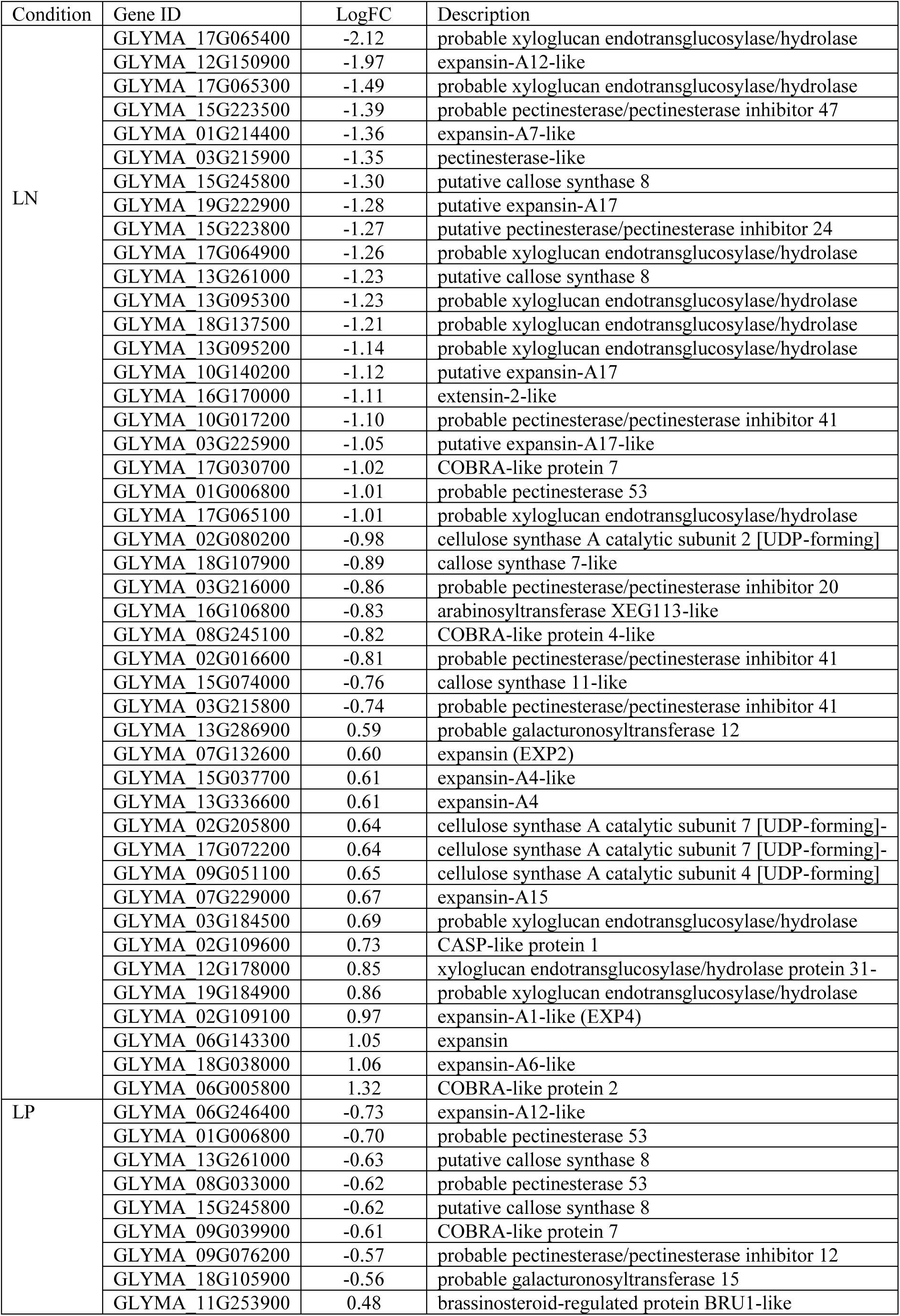
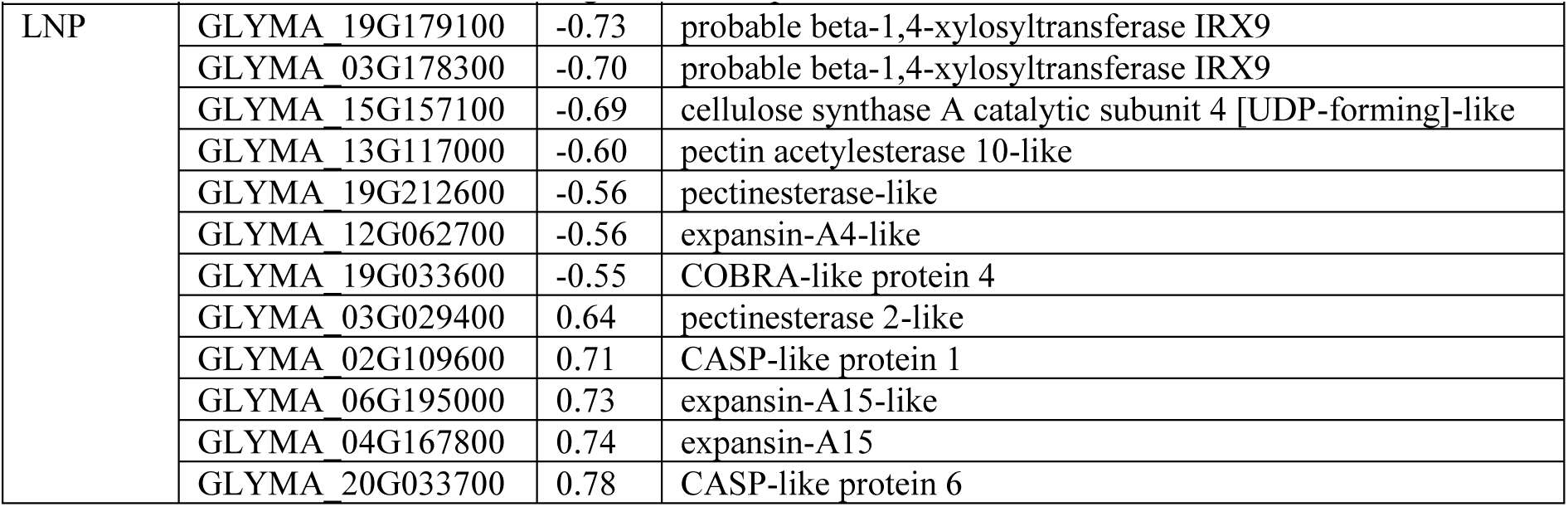
Log foldchanges of DEGs in plant-type cell wall organization in LN, LP, and LNP conditions

**Supplemental Table S5.**
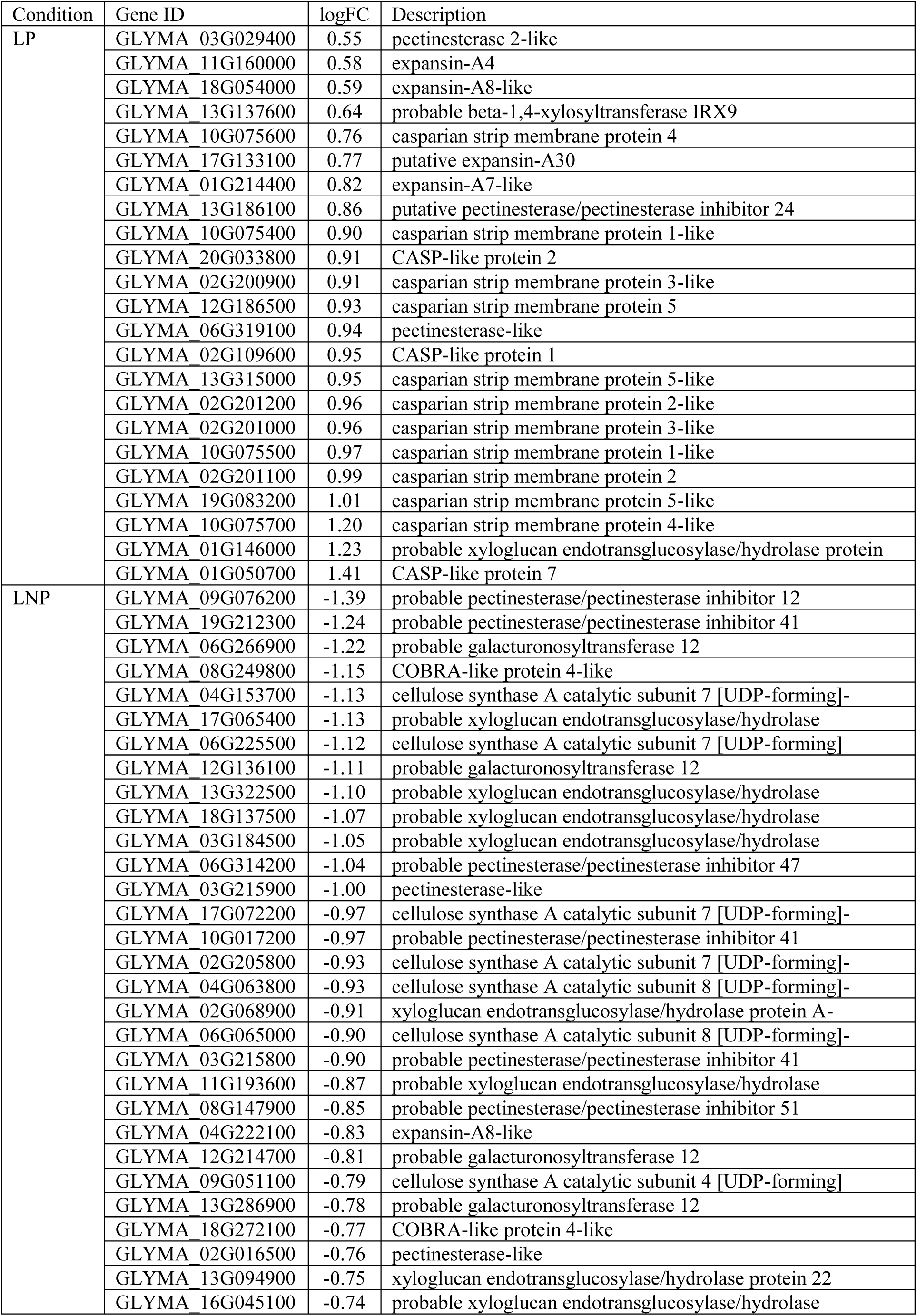

**Supplemental Figure S1.**
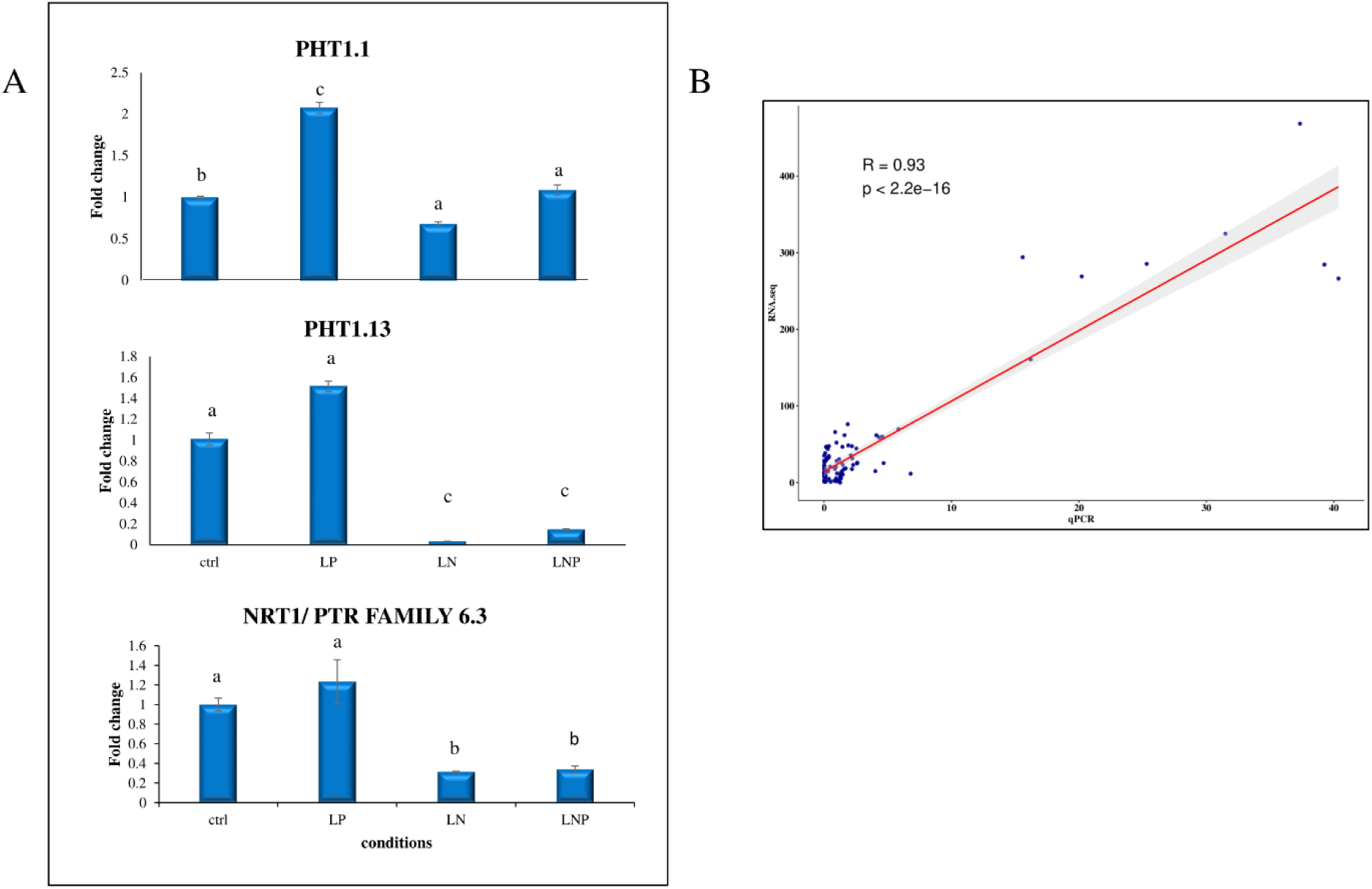
RT-qPCR validation of RNA-seq data. A. Transporter genes for nitrate (NRT1/PTR family 6.3) and phosphate transporter (PHT1.1 and PHT1.13) were used for RT-qPCR analysis using 3 biological and 2 technical replicates. The data are shown as means ± SD, the values in controls were set to 1. The significant differences (Tukey test <0.05) are indicated with asterisks with different superscript letters.. B. Correlation plot of RPKM data from RNA-seq and relative expressions from RT-qPCR

**Supplemental Figure S2.**
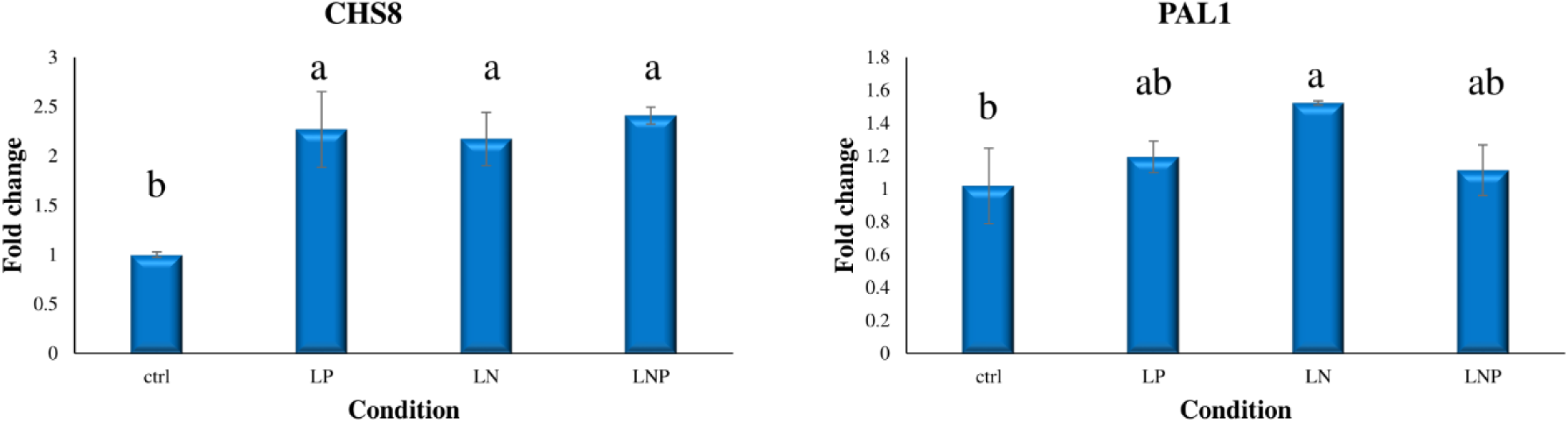
Gene expression analysis of some flavonoid genes in response to nitrogen (LN), phosphorus (LP), and the combined deficiency (LNP). Expression levels were determined by RT-qPCR using 3 biological and 2 technical replicates. The data are shown as means ± SD, the values in controls were set to 1. The significant differences (Tukey test <0.05) are indicated with asterisks with different superscript letters.

